# Parasitoid wasp venom targets host immune cell production in a *Drosophila*-parasitoid interaction

**DOI:** 10.1101/2020.12.01.406736

**Authors:** Jordann E. Trainor, KR Pooja, Nathan T. Mortimer

## Abstract

The interactions between *Drosophila melanogaster* and the parasitoid wasps that infect *Drosophila* species provide an important model for understanding host-parasite relationships. Following parasitoid infection, *D. melanogaster* larvae mount a response in which immune cells (hemocytes) form a capsule around the wasp egg, which then melanizes leading to death of the parasitoid. Previous studies have found that host hemocyte load, the number of hemocytes available for the encapsulation response, and the production of lamellocytes, an infection induced hemocyte type, are major determinants of host resistance. Parasitoids have evolved various virulence mechanisms to overcome the immune response of the *D. melanogaster* host, including both active immune suppression by venom proteins and passive immune evasive mechanisms. We find that a previously undescribed parasitoid species, *Asobara sp. AsDen*, utilizes an active virulence mechanism to infect *D. melanogaster* hosts. *Asobara sp. AsDen* infection inhibits host hemocyte expression of *msn*, a member of the JNK signaling pathway, which plays a role in lamellocyte production. *Asobara sp. AsDen* infection restricts the production of lamellocytes as assayed by hemocyte cell morphology and altered *msn* expression. Our findings suggest that *Asobara sp. AsDen* venom targets host signaling to suppress immunity.

**Declarations:** *Funding:* This work was supported by the National Institute of General Medical Sciences of the National Institutes of Health under Award Number R35GM133760.

*Availability of data and material:* Sequence data has been deposited in GenBank under accession # MT498809. Custom BLAST databases are available on request to corresponding author.

*Authors’ contributions:* Conceived of or designed study: J.E.T., N.T.M.; Performed research: J.E.T., P.K.; Analyzed data: J.E.T., P.K., N.T.M.; Wrote the paper: J.E.T., P.K., N.T.M.

## 1. Introduction

Parasitoid wasps that infect *Drosophila* are a valuable model for understanding parasite behaviour and have provided important ecological and molecular insights into host-parasite interactions [1–3]. In this system, parasitoids infect larval *Drosophila* and following infection, *Drosophila* mount a cellular encapsulation response to overcome the invader [4]. This encapsulation response is highly conserved among arthropods [5–9], and encapsulation ability is an important determinant of pathogen resistance in insect vectors of human disease [10–12]. The encapsulation response in *Drosophila melanogaster* is mediated by hemocytes (immune cells), including circulating macrophage-like cells known as plasmatocytes, and lamellocytes, a highly specialized infection-induced immune cell subtype [13]. Plasmatocytes are physiologically activated by parasitoid wasp infection, and following activation they migrate and adhere to the surface of the parasitoid egg [14, 15]. Immune stimulation also triggers the production of lamellocytes [16, 17], which adhere to the plasmatocyte cell layer and form a melanized capsule around the egg, killing the developing parasitoid [15, 18]. There are multiple routes for lamellocyte production, including the transdifferentiation of plasmatocytes in circulation or within sessile populations, and differentiation directly from prohemocyte precursors in the lymph gland, the main hematopoietic organ in *Drosophila* [19–21].

It has been proposed that the main determinant of *Drosophila* immune resistance to parasitoid infection is host hemocyte load [22]. In this context hemocyte load refers both to the number and activity of hemocytes found in circulation and the potential for the production of additional hemocytes following infection. Studies have found that an increased number of hemocytes confers resistance to parasitoid infection in *D. melanogaster* and other *Drosophila* species [23–27], and that the production and function of lamellocytes is critical for a successful encapsulation response [18, 22, 27].

*Drosophila* parasitoid wasps have evolved multiple mechanisms that allow them to evade or overcome the host immune response, the most prevalent of which is the transfer of venom virulence proteins into the host during infection. Because of the importance of hemocyte number for resistance, many of these parasitoid virulence mechanisms target host hemocytes. This includes venom virulence proteins that act on host hemocytes in a variety of ways including inducing hemocyte lysis [28], promoting death of hemocyte precursor cells [29, 30], and inhibition of hemocyte function leading to immunodeficiency [14, 18, 31–34]. Many of these venom proteins specifically target lamellocytes [17, 18, 28, 34, 35], reinforcing the vital role that this hemocyte subtype plays in the encapsulation response. The outcome of these venom activities is to suppress host hemocyte load either by reducing the number or function of these immune cells.

Along with these active immune suppression mechanisms, parasitoids can also use passive immune evasive mechanisms to escape encapsulation [36, 37]. Proposed passive mechanisms include the ability of parasitoid eggs to bind to host tissues as a form of camouflage from the immune response [14, 36, 38], an increase in parasitoid egg size following infection [39, 40] or superparasitism, where a single host is multiply infected by conspecific parasitoids, which has been suggested to increase parasitoid infection success [40–43].

The conservation of the encapsulation response in human disease vectors, and the use of parasitoid wasps as biological control agents makes understanding parasitoid virulence strategies an important research goal. In the present study, we describe an uncharacterized parasitoid species of the genus *Asobara* that utilizes a venom mediated mechanism to suppress *D. melanogaster* lamellocyte development and thereby overcome host immune defense.

## 2. Results

### 2.1. AsDen is a strain of an undescribed Asobara species

Female Braconid wasps were caught in Denver, CO, USA and allowed to infect the *OstΔ*^*EY02442*^ encapsulation deficient *D. melanogaster* strain [18]. These infections resulted in an all-female parthenogenetic strain which was reared in the lab for several generations prior to beginning experimentation. We sequenced the COI gene from this wasp strain and compared the sequence to COI sequences from known Braconid species. Our sequence analysis suggests that the strain is a previously undescribed species of the genus *Asobara*. We will refer to this wasp species using the name *Asobara sp. AsDen* or by the strain name *AsDen* to indicate the genus and location of collection.

Our BLAST analysis of *Asobara sp. AsDen* reveals that the most closely related species are additional uncharacterized species of *Asobara* identified in recent efforts to catalog arthropod biodiversity (Table 1) [44–46]. In order to further characterize the evolutionary relationships between *Asobara sp. AsDen* and these other species, we performed phylogenetic analysis using COI sequences. We find that *Asobara sp. AsDen* forms a supported clade with the species *Asobara sp. ABZ3773* and *Asobara sp. ABX5347* [46] (Figure 1A). Interestingly these species are also found in North America (Table S1), further suggesting a recent evolutionary relationship. Additional phylogenetic analysis with previously studied species of *Asobara* suggests that the species group including *Asobara sp. AsDen, Asobara sp. ABZ3773* and *Asobara sp. ABX5347* is most closely related to *Asobara triangulata*, a species known from a single sample collected in Yunnan, China [47], *Asobara mesocauda*, a species collected in South Korea and China [47], and the well-studied species *Asobara rufescens* and *Asobara tabida* which have both been found in Asia, Europe and North America [46–48] (Figure 1B and Table S2).

**Table 1.** BLAST results comparing the *AsDen* COI DNA sequence against a custom database of 353 *Asobara* COI sequences. The species name, sequence accession number, score (bits) and identity (%) for the top scoring hits by species are displayed.

| Species Designation | Accession # | Score (Bits) | Identity (%) |
| --- | --- | --- | --- |
| *Asobara* sp. ABZ3773 | KR886087.1 | 974 | 94 |
| *Asobara* sp. ABX5347 | JN293161.1 | 924 | 93 |
| *Asobara* sp. ACF3746 | HQ929638.1 | 913 | 92 |
| *Asobara* sp. ACE4721 | JN293665.1 | 907 | 92 |
| *Asobara* sp. ACR5030 | MF936732.1 | 902 | 92 |
| *Asobara* sp. ACF3747 | HQ930298.1 | 896 | 92 |
| *Asobara* sp. AAE0947 | HQ106668.1 | 891 | 92 |

**Figure 1.**
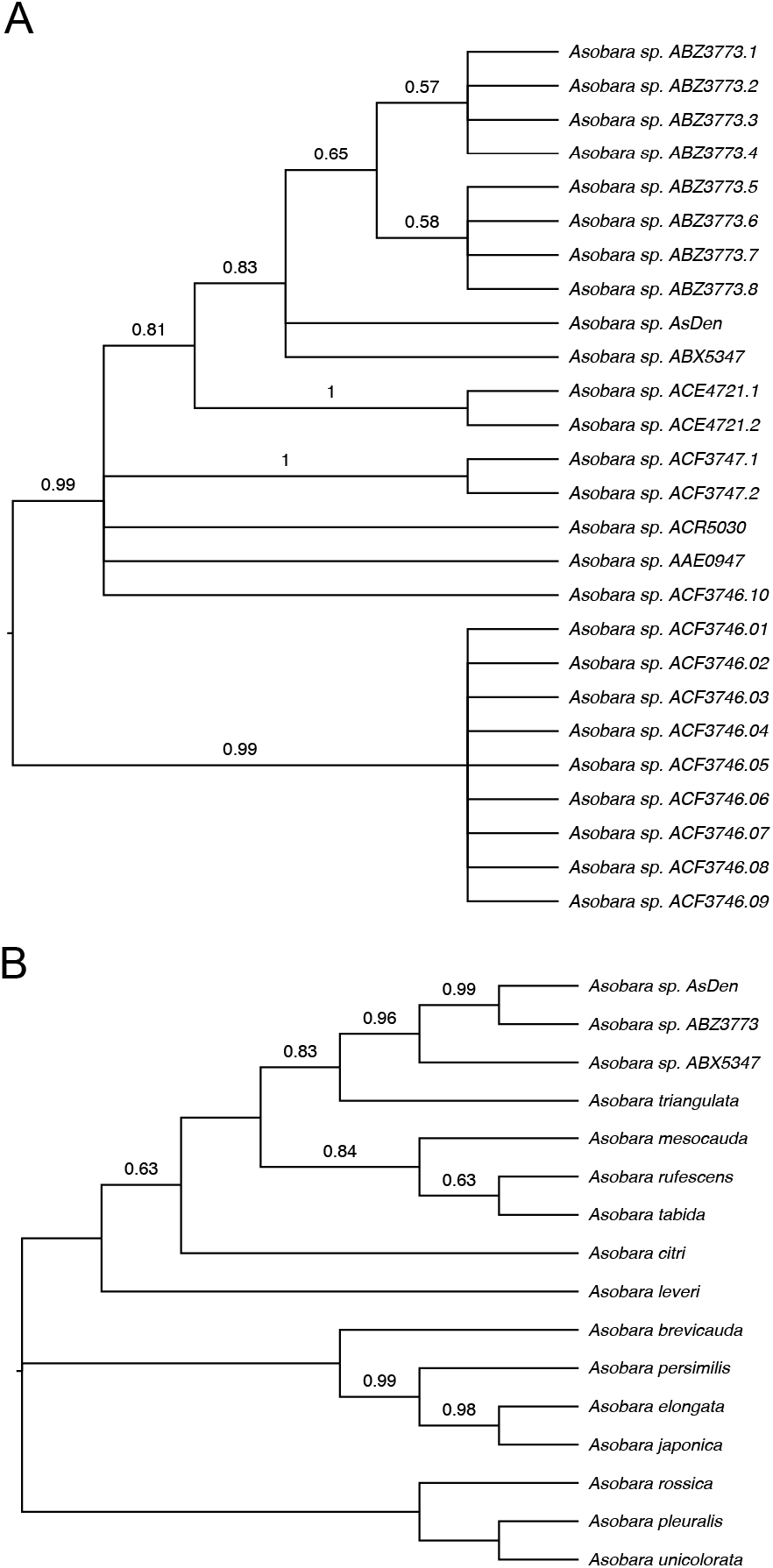
(A-B) Phylogenetic analysis of the COI gene in *Asobara sp. AsDen* with other species of the genus *Asobara*. The evolutionary history was inferred by using the Maximum Likelihood method and the tree with the highest log likelihood is shown. The proportion of trees from 1000 bootstrap replicates in which the associated taxa clustered together is displayed, and values below 0.5 are not shown. (A) Phylogeny of *Asobara sp. AsDen* with sequences from 25 individuals belonging to closely related undescribed *Asobara* species (see Supplemental Table 1 for sequence information). Strains of the same species have a numerical suffix appended to the species name. (B) Phylogeny of *Asobara sp. AsDen* with sequences from well-studied species of *Asobara* (see Supplemental Table 2 for sequence information).

### 2.2. Asobara sp. AsDen avoids encapsulation by D. melanogaster hosts

*AsDen* wasps readily infect *D. melanogaster* larvae, with 98.8% of hosts infected after a 72-hour exposure period (n = 90 larvae). We find that the *D. melanogaster* immune response successfully encapsulated only 36.6% of *AsDen* eggs (n = 372 eggs), and that 34.8% of infected *D. melanogaster* larvae were able to encapsulate all of the infecting *AsDen* eggs (n = 89 infected larvae). To survive infection, a host must encapsulate every infecting parasitoid egg, so these data suggest a high rate of successful parasitization of *D. melanogaster* hosts by *AsDen*. Interestingly, 77.5% of infected *D. melanogaster* larvae were infected more than once during the exposure period, for an average of 4.2 eggs/infected host larva (n = 89 infected larvae). We find a significant negative correlation between the number of eggs laid per larva and the proportion of eggs that are encapsulated (Figure 2A; Pearson’s r = −0.576, p < 0.001). Taken together, these data suggest that *AsDen* can successfully parasitize *D. melanogaster* hosts and that multiply infected host larvae are less likely to survive infection.

**Table 2.** Eigenvalues and factor loading for the first two dimensions (PC1 and PC2) from PCA of cell morphology and fluorescence intensity of all hemocytes extracted from *L. boulardi* and *AsDen* infected *msn-mCherry* larvae, as shown in Figure 4A-B.

| Variable | PC1 | PC2 |
| --- | --- | --- |
| Eigenvalue | 1.624 | 0.867 |
| Variance (%) | 54.13 | 28.90 |
| *Factor Loading* |  |  |
| Cell Size | 0.647 | -0.262 |
| Cell Circularity | 0.636 | -0.332 |
| Fluorescence intensity | 0.420 | 0.906 |

**Figure 2.**
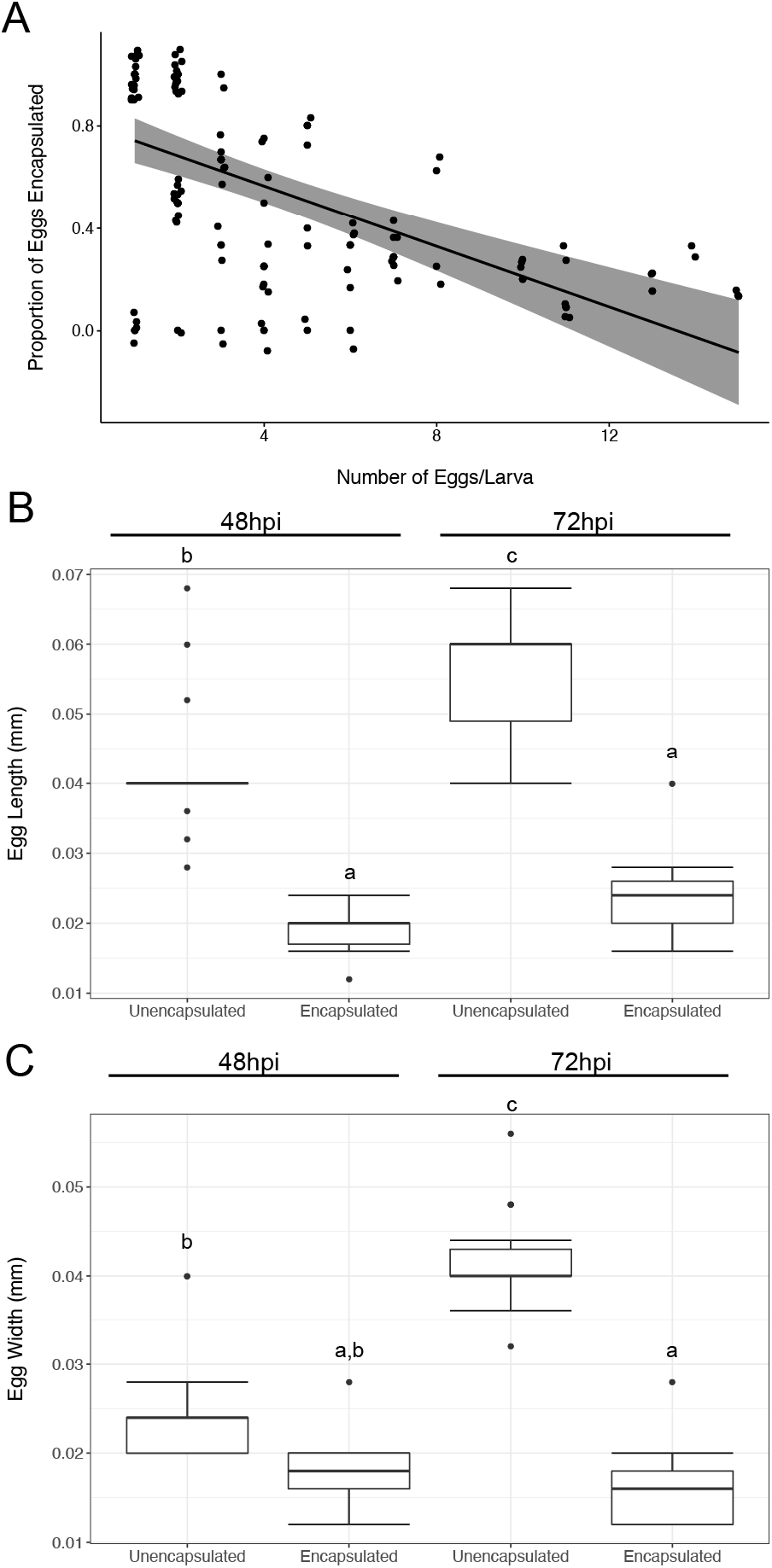
(A) Scatterplot showing the correlation between the number of infections and the proportion of *AsDen* eggs encapsulated in *w*^*1118*^ hosts. Individual data points are shown, and the 95% confidence interval is shaded in grey. The length (B) and width (C) of both unencapsulated and melanotically encapsulated *AsDen* eggs were determined at 48- and 72-hours post infection (hpi). Data are displayed as box plots, with calculated outlier data shown as individual points. Letters (a-c) indicate significance groups within each experiment as determined by Tukey’s HSD.

Similar to other *Asobara* species [39, 40], we find that *AsDen* eggs continue to grow in size as they develop in *D. melanogaster* hosts. Eggs were dissected from infected *D. melanogaster* larvae at 48hpi and 72hpi and the length and width of each individual egg was determined. Unencapsulated eggs continue to increase in length (Figure 2B; t = 4.309, p < 0.001) and width (Figure 2C; t = 7.678, p < 0.001) between 48hpi and 72hpi. To verify that encapsulation was arresting parasitoid development, the length and width of individual encapsulated and melanized eggs were determined at 48hpi and 72hpi to compare with unencapsulated eggs. We find that the melanized eggs are significantly shorter (Figure 2B; t = −8.285, p < 0.001), and narrower (Figure 2C; t = −8.382, p < 0.001) than unencapsulated eggs at 72hpi. Additionally, the increase in size that is seen in unencapsulated eggs is arrested in encapsulated eggs, with no significant size differences observed in encapsulated eggs dissected at 48hpi and 72hpi (Figure 2B-C; length: t = 1.237, p = 0.602; width: t = −0.624, p = 0.9221).

### 2.3. Host lamellocyte production is impaired in Asobara sp. AsDen infected larvae

Many parasitoid species transfer venom virulence proteins to their host during infection to suppress hemocyte number or activity. Often these virulence proteins target lamellocytes, a parasitoid-infection induced hemocyte subtype that is required for a successful encapsulation response [27–30]. Lamellocytes are larger and less circular than other hemocytes, and can be distinguished from other hemocyte subtypes both by their unique morphology and by the specific expression of *misshapen* (*msn*) [49, 50]. Lamellocytes are produced both by the direct differentiation of prohemocytes in the hematopoietic lymph gland and by the transdifferentiation of circulating or sessile plasmatocytes [16, 19–21], and both routes result in *msn* expression [49].

To assay the production of lamellocytes in *AsDen* infected larvae, we used a fluorescent cytometer to take high throughput measurements of cell size, cell perimeter, cell circularity, and *mCherry* fluorescence intensity from hemocytes isolated from infected larvae of the *msn-mCherry* strain. This strain expresses *mCherry* as a fluorescent reporter of *msn* expression [51]. Because lamellocyte production is induced by parasitoid infection, *msn* is not expressed in the hemocytes of naïve larvae, and so we used infection with the parasitoid *Leptopilina boulardi* as a comparison for *AsDen* venom activity. *L. boulardi* infection does not inhibit expression of *msn* or lamellocyte development and so provides a reliable control [17, 52, 53]. We find that following *L. boulardi* infection, 45.1 ± 4.2% of circulating hemocytes express the *msn-mCherry* reporter (n = 32,176 hemocytes). In these *L. boulardi* infected larvae, the *msn-mCherry* positive cells are larger (cell size: t = 29.265, p < 0.001; cell perimeter: t = 29.442, p < 0.001) and less circular (t = 21.796, p < 0.001) than cells not expressing *msn-mCherry*, consistent with the described properties of lamellocytes [50].

We find that 21.4 ± 1.8% of hemocytes in *AsDen* infected *msn-mCherry* larvae were *msn* positive (n = 53,908 hemocytes), a significantly lower proportion than observed in stage matched *L. boulardi* infected *msn-mCherry* larvae (Figure 3A; z = 7.328, p < 0.001). We further find that among the *mCherry* positive hemocytes, cells from *AsDen* infected larvae had significantly lower fluorescence intensity compared to cells from *L. boulardi* infected larvae (Figure 3B; z = 4.838, p < 0.001). These differences in *msn* expression may be predicted to result in differences in hemocyte morphology from *L. boulardi* and *AsDen* infected larvae. To better compare cell morphology between infections, we used principal components analysis (PCA) to reduce the cell size, cell perimeter and cell circularity measures from the cytometer data to a single dimension. The first principal component of this cell morphology PCA (PCM) has an eigenvalue of 2.35 and explains 78.4% of the variance among these data, suggesting that it accurately captures the data describing hemocyte morphology. We find that PCM values differ significantly between hemocytes from *AsDen* and *L. boulardi* infected larvae (Figure 3C; t = 17.03, p < 0.001), implying that hemocyte morphology does vary by infection condition.

**Figure 3.**
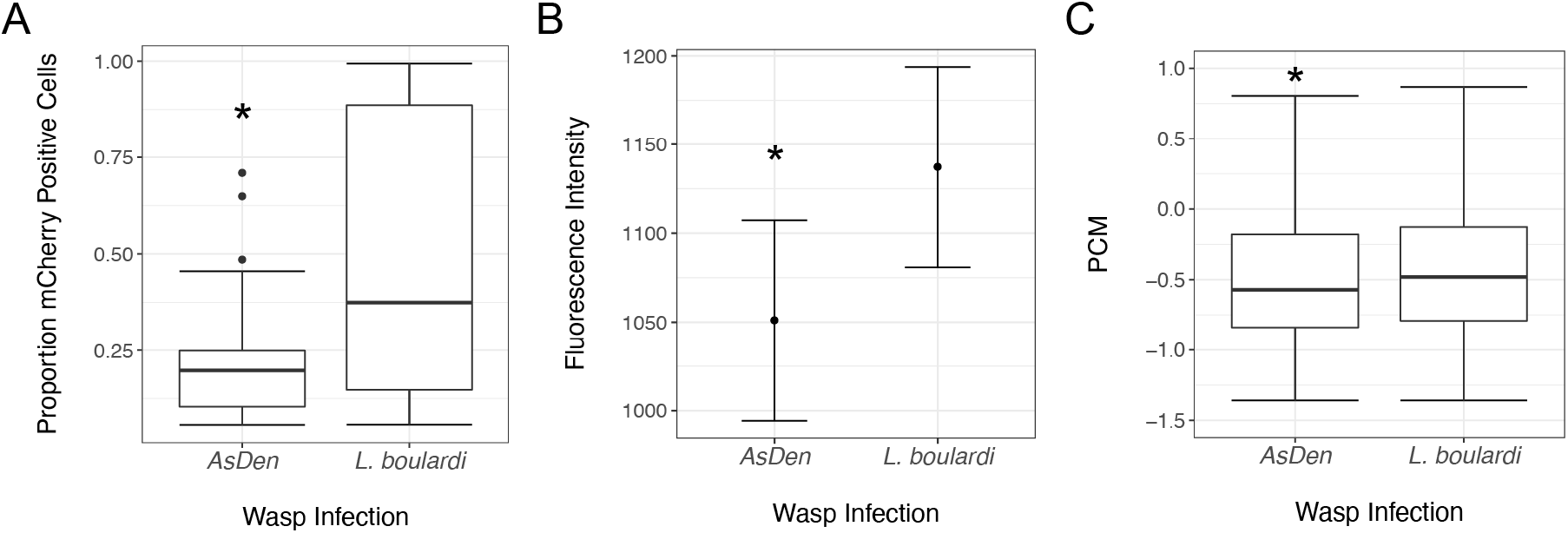
(A) The proportion of hemocytes positive for *mCherry* isolated from *AsDen* and *L. boulardi* infected *msn-mCherry* larvae 72 hpi. Data are displayed as box plots, with calculated outlier data shown as individual points. (B) Calculated fluorescence intensity of *mCherry* positive hemocytes isolated from *AsDen* and *L. boulardi* infected larvae 72 hpi. Data are displayed as the mean fit (point) of the effect of parasitoid species on fluorescence intensity ± standard error. (C) PCM values calculated from hemocytes isolated from *AsDen* and *L. boulardi* infected larvae 72 hpi. Data are displayed as box plots. * indicates p < 0.05 compared to *L. boulardi* infected larvae.

To further characterize the hemocyte populations in *AsDen* and *L. boulardi* infected larvae, we performed a second PCA using the previously listed cell morphology features and *mCherry* fluorescence intensity data. We plotted the first two dimensions of this PCA (PC1 and PC2; Table 2), and we find that hemocytes from *L. boulardi* infected larvae (red triangles in Figure 4A-B) largely cluster into two groups, distinguished by morphology and fluorescence intensity. Although hemocytes from *AsDen* infected larvae fall into a similar pattern (black circles in Figure 4A-B), one of these groups is greatly reduced. The same pattern is replicated when only data from *mCherry* positive cells are used for the PCA (Figure 4C-D). However, the PCA plots derived from *mCherry* negative hemocyte properties are indistinguishable between *AsDen* and *L. boulardi* infected larvae (Figure 4E-F). These data support the hypothesis that *msn* expressing hemocytes are differentially affected by the parasitoid infections. Based on the role of *msn* in lamellocyte production and the observed morphology differences, these data suggest that lamellocyte production is impaired following *AsDen* infection.

**Figure 4.**
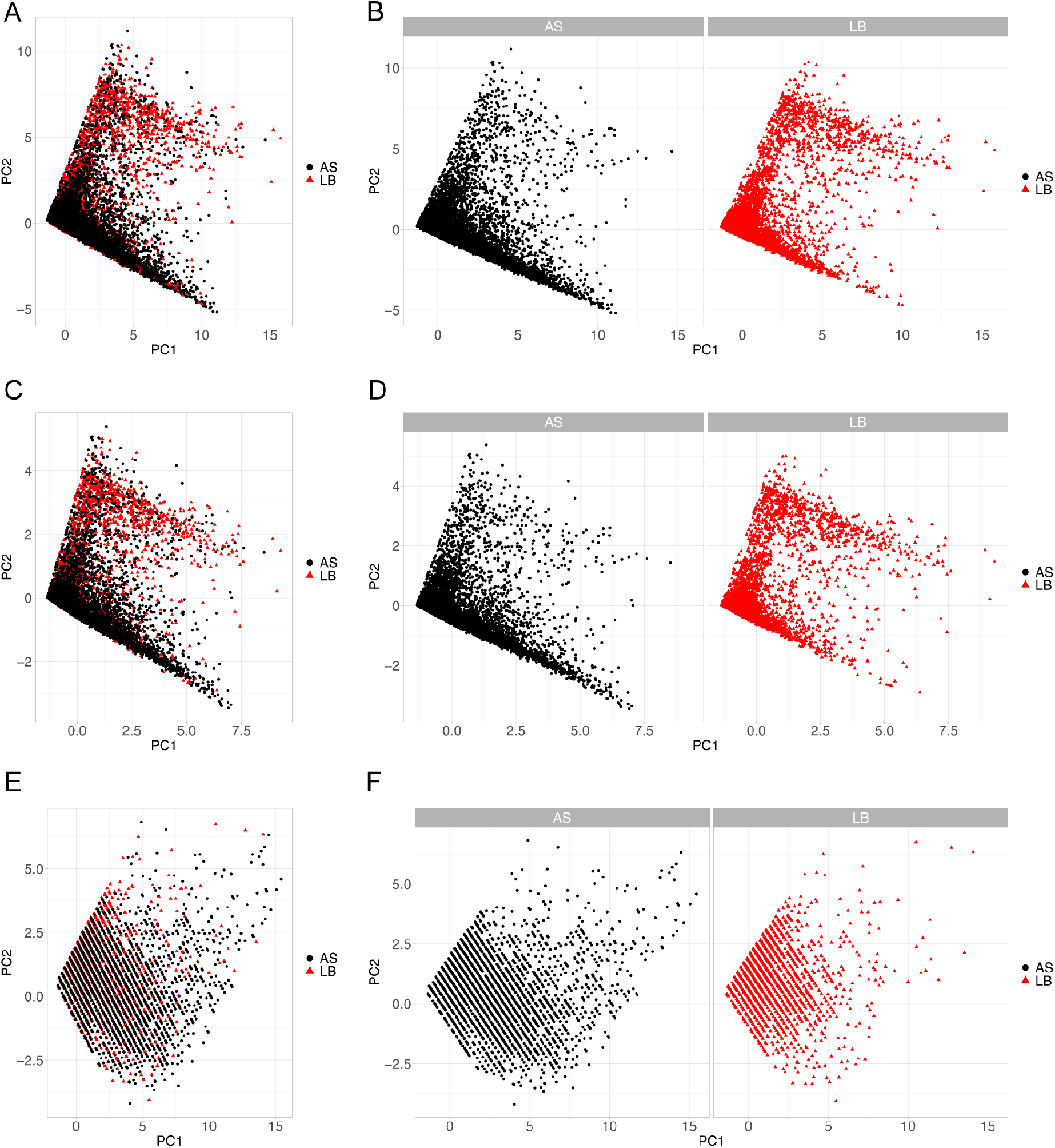
Plots of the first two principal components from a PCA of cell morphology and fluorescence intensity performed on (A-B) all hemocytes, (C-D) *msn-mCherry* positive hemocytes and (E-F) *msn-mCherry* negative hemocytes. Hemocytes were extracted 72hpi from *msn-mCherry* larvae infected by the indicated parasitoid. Hemocytes from *AsDen* infected larvae are shown as black circles and as the left panel of faceted images (B,D,F) and hemocytes from *L. boulardi* infected larvae are shown as red triangles and as the right panel of faceted images.

## 3. Discussion

Our findings suggest that a previously uncharacterized parasitoid species from the genus *Asobara*, represented here by the *AsDen* strain, can successfully parasitize *D. melanogaster*. *Asobara sp. AsDen* is evolutionarily related to other *Drosophila* infecting parasitoids including *A. tabida*, although the host ranges of the more closely related, uncharacterized species found in North America are unknown. To characterize the effects of *AsDen* infection on host hemocyte load, and specifically hemocyte morphology and *msn* expression, we compared the properties of hemocytes from *AsDen* infected hosts to hemocytes from *L. boulardi* infected hosts. *L. boulardi* infection triggers *msn* expression and lamellocyte production, and *L. boulardi* venom has no known impact on these processes [17, 38, 52, 53], suggesting that this infection can serve as a useful control for our analyses.

We find that *AsDen* infection has a distinct effect on both hemocyte morphology and *msn* expression in host hemocytes when compared with *L. boulardi* infection. In the encapsulation response, *msn* is expressed in lamellocytes following infection and *msn* signaling activity is required for lamellocyte production [51, 54]. The proportion of *msn*-positive immune cells is lowered following *AsDen* infection, and *msn* expression levels are decreased in immune cells isolated from *AsDen* infected larvae in comparison with *L. boulardi* infected larvae (Figure 3). These findings suggest that *AsDen* infection inhibits host immune signaling leading to the failure to properly promote lamellocyte specification or development. In agreement with this hypothesis, we find that while hemocytes from *L. boulardi* infected hosts cluster into two populations based on their morphology and *msn* expression levels, one of these populations is greatly reduced in *AsDen* infected hosts (Figure 4A-B). Lamellocytes tend to be larger and more irregularly shaped than plasmatocytes [50]. An examination of the factor loading from our cell morphology and fluorescence intensity PCA results (Table 2), suggests that the reduced cell population in *AsDen* infected larvae tends to be larger, less circular and *msn* positive (Figure 4), all of which are consistent with a specific deficit in lamellocyte production. The finding that this alteration in hemocyte characteristics in observed in *msn* positive cells (Figure 4C-D) but not *msn* negative cells (Figure 4E-F) further suggests that the activity of *AsDen* venom is specifically targeting *msn* and/or lamellocyte production.

*msn* is a member of the JNK signal transduction pathway [55] and *msn-mCherry* provides a readout of JNK pathway activity [51]. This suggests that the JNK signaling pathway may be inhibited in *AsDen* infected larvae. We have yet to determine the molecular mechanism underlying JNK inhibition in *AsDen* infected larvae, but we propose it could act either directly through inhibiting one or more components of the JNK pathway or indirectly by blocking upstream pathway activation to inhibit lamellocyte production. The JNK pathway plays a conserved role in immunity in *Drosophila* and a wide range of species [9, 56, 57]. In *D. melanogaster*, genes in the JNK pathway are associated with resistance to parasitoids [58, 59], and are required for lamellocyte production in response to infection [51]. To our knowledge, *AsDen* is the first *Drosophila* parasitoid suggested to inhibit JNK signaling, however the JNK pathway is targeted by a wide range of other pathogens in variety of hosts [60–62].

It is notable that *AsDen* infected larvae do still produce *msn* positive hemocytes, suggesting that lamellocyte differentiation and JNK signaling are not completely abolished. Additionally, even though the morphological changes leading to lamellocyte production are impaired in *AsDen* infected larvae, the cell morphology of *msn* expressing hemocytes is different from non-*msn* expressing hemocytes. These data suggest that *AsDen* venom may be inhibiting a specific aspect of lamellocyte transdifferentiation or maturation, consistent with the finding that *msn* expression coincides with early morphological changes in transdifferentiating hemocytes [49]. Recent studies have uncovered a broader range of *Drosophila* hemocyte subtypes than previously appreciated [63–66], and future investigation into this complexity may help to unravel the specific effects of *AsDen* venom on host hemocytes and lamellocyte production.

Along with restricted lamellocyte production, *AsDen* infected hosts have a limited encapsulation response. Interestingly, we find a negative correlation between the number of times a host larva has been infected and its encapsulation ability (Figure 2A). Multiple infections of a single host by conspecific parasitoids is known as superparasitism [67], and is commonly observed across many parasitoid species both in laboratory conditions and in nature. The negative effect of superparasitism on host resistance observed in our study may be due the additive effects of multiple envenomations on host lamellocyte production; perhaps additional “doses” of venom are able to more completely suppress lamellocyte production. However, we cannot rule out the possibility that superparasitism is acting through an alternative mechanism such as passive immune evasion [36, 37]. Supernumerary infections by the parasitoids *Pseudapanteles dignus* and *A. tabida* have been shown to increase the likelihood of successful parasitization [40, 42], suggesting that superparasitism itself may contribute to the ability of the parasitoid egg to escape from encapsulation. Parasitoids generally avoid superparasitism; most parasitoid species are able to perceive the presence of eggs from a conspecific female [39, 41, 68], and in previous work, we found that using the identical experimental set up with other parasitoid species consistently yields average infection rates of 1-1.2 eggs per infected larva [14, 38], in contrast to the 4.2 eggs per infected larva observed for *AsDen*. Many known instances of superparasitism are driven by external factors such viral infections [69–71], but this has not yet been determined in this case.

In *Asobara sp. AsDen* and many other parasitoid species, virulence appears to be largely driven by a single strategy, for example the passive immune avoidance of *A. tabida* or the immune suppressive venoms of *AsDen*, *Asobara citri*, *Asobara japonica* or various species of Figitid parasitoid wasps [30, 32, 36, 72–74]. However, both *L. boulardi* and *Ganaspis hookeri* appear to use a combined strategy of venom-mediated immune suppression and passive avoidance [14, 37, 38], suggesting that further study may uncover more complex virulence strategies across a range of parasitoids than previously appreciated. Further, while *A. tabida* is the most closely related of the well-studied parasitoid species to *Asobara sp. AsDen*, its venom has been shown to cause paralysis and inhibit host development with only limited immune-suppressive effects [36, 75–79]. This is not entirely unexpected, as other closely related parasitoid species have distinct virulence strategies and venom composition [37, 38, 80]. It has also been demonstrated that different strains of a single parasitoid species can possess different virulence activities [81–83]. As *AsDen* is the only known strain of its species, we aren’t able to determine how conserved this activity may be with other strains, although this will hopefully be investigated as more strains of this species are identified.

Our findings support the idea that overcoming host hemocyte load is a critical determinant of parasitization success for parasitoid wasps of *Drosophila*. Since *Drosophila* are a valuable model for understanding the immune defenses of insect vectors of human disease and agricultural pests, these findings may provide insight into the interactions between insect vectors and invading pathogens and may have implications for the selection and use of parasitoid wasps in biological control applications.

## 4. Materials and Methods

### 4.1. Insect Strains

Two females from an unknown Braconid parasitoid wasp species were collected from a fruit trap in Denver, Colorado, USA and were maintained on the encapsulation deficient *D. melanogaster* mutant strain *OstΔ*^*EY02442*^ (BDSC: 15565) [18] from the Bloomington Drosophila Stock Center. A sub-strain was established from a single parthenogenetic foundress and will be referred to as *AsDen*. The study also uses the parasitoid wasp *Leptopilina boulardi* (strain Lb17) [38] which is maintained in the laboratory on the *Canton S D. melanogaster* strain. The following additional *D. melanogaster* strains were used in this study: *w*^*1118*^ (BDSC: 5905) from the Bloomington Drosophila Stock Center; and *msn-mCherry* [51], provided by Dr. Robert Schulz.

### 4.2. Parasitoid species determination

Genomic DNA was extracted from *AsDen* using standard methods. The COI gene was amplified using the “Folmer” primers [84] LCO1490 (primer sequence: GGTCAACAAATCATAAAGATATTGG) and HCO2198 (primer sequence: TAAACTTCAGGGTGACCAAAAAATCA), and sequenced at the UIUC Core Sequencing Facility (Urbana, IL). The resulting Sanger sequencing reads were aligned using 4Peaks software (A. Griekspoor and Tom Groothuis, nucleobytes.com). The *Asobara sp. AsDen* COI DNA sequence was submitted to GenBank (accession # MT498809) The resulting DNA sequence was compared against all Hymenopteran sequences using the Basic Local Alignment Search Tool (BLAST) available through the National Center for Biotechnology Information (NCBI) [85]. For further sequence analysis, we constructed a custom BLAST database of all 353 *Asobara* COI sequences available from NCBI (accessed April 11, 2020) using BLAST+ (version 2.5.0) [86]. This custom BLAST database is available upon request.

### 4.3. Phylogenetics

Phylogenetic analyses were conducted in MEGA X [87, 88] using COI DNA sequences. For the first analysis, *AsDen* was compared to the 25 most highly homologous *Asobara* sequences as determined by BLAST+ (Supplemental Table 1) [44–46]. For the second analysis, the species group including *AsDen* found in the first analysis was compared against 13 well-studied species of *Asobara* (Supplemental Table 2) [26, 47, 89, 90]. For both analyses, the evolutionary history was inferred by using the Maximum Likelihood method and Kimura 2-parameter model with 1000 bootstrap replicates [91]. The initial tree for the heuristic search was obtained automatically by applying Neighbor-Join and BioNJ algorithms to a matrix of pairwise distances estimated using the Maximum Composite Likelihood (MCL) approach in MEGA X, and then selecting the topology with superior log likelihood value. Branches corresponding to partitions reproduced in less than 50% of the bootstrap replicates were collapsed. All positions containing gaps and missing data were eliminated. The resulting phylogenetic trees were visualized using FigTree (version 1.4.3, http://tree.bio.ed.ac.uk/).

### 4.4. Parasitoid infection

For infection with parasitoid wasps, 30 late second instar larvae from the *w*^*1118*^ strain were placed on 35mm Petri dishes filled with Drosophila medium together with 3 *AsDen* wasps at 25°C. Larvae were dissected at 48- or 72-hours post infection (hpi) as noted. The infected larvae were then scored for the total number of parasitoid eggs and the numbers of encapsulated and non-encapsulated eggs. For size experiments, the length and width of each egg was determined using an E-series Reticle (Leica Microsystems). Egg length was measured from pole to pole and egg width was measured across the widest region perpendicular to the length axis. All experiments were performed in triplicate.

### 4.5. msn expression and cell morphology analyses

The *msn-mCherry D. melanogaster* strain was used to assay expression of *msn*. This strain carries a transgenic construct containing the *msn-F9* enhancer upstream of the *mCherry* red fluorescent protein [51]. Second instar *msn-mCherry* larvae were infected by either *AsDen* or *L. boulardi* as described above, with three biological replicates for each infection condition. Host hemocytes were isolated 72hpi and added to a Tali Cellular Analysis Slide (Invitrogen). Hemocytes were allowed to adhere for 30 minutes and then cell number, size, perimeter, circularity and red fluorescence intensity were measured using a Tali Image-Based Cytometer (Invitrogen). For each replicate, we imaged 20 fields of cells, with an average of 717.4 cells per field, and a range of 194 to 1455 cells for a total of 32,176 hemocytes from *L. boulardi* infected larvae and 53,908 hemocytes from *AsDen* infected larvae. Cytometer data were filtered to only include single cells using the Tali software count function and size-gating, prior to further analysis.

### 4.6. Data analysis

All statistical analyses were done in the R statistical computing environment [92] using the multcomp [93], lme4 [94], lmerTest [95], plyr [96], FactoMineR [97], factoextra [98] and ggplot2 [99] packages. Analysis of Variance (ANOVA) was used to test the relationship between egg size and time or encapsulation status. Tukey’s Honest significant difference (HSD) test was used for multiple comparisons of egg size. Pearson’s product-moment correlation was used to test for correlations between egg number and encapsulation status. Mixed linear models, with replicate as a random effect, were used to test for differences in *msn*-*mCherry* fluorescence intensity and proportion of *mCherry* positive cells between *AsDen* and *L. boulardi* infections. Welch Two Sample t-tests were used to compare immune cell morphology data between *AsDen* and *L. boulardi* infections.

To characterize hemocyte populations, we used PCA on the red fluorescence intensity, cell size, cell perimeter and cell circularity measures from the cytometer data. A circularity value of 1.0 is considered perfectly circular, and values either greater or less than 1.0 are increasingly less circular. To account for this, circularity values were log2 transformed and the absolute value of these transformed values were used for PCA. Other measures were used for PCA without transformation. This analysis was repeated separately on gated fluorescence data, generating distinct PCA scores for *mCherry* positive hemocytes and *mCherry* negative hemocytes.

## Acknowledgements

The authors would like to thank Dr. Robert Schulz for providing the *msn-mCherry* line. We would also like to thank members of the Mortimer Cellular Immunology lab for their input throughout the project. Stocks obtained from the Bloomington Drosophila Stock Center (NIH P40OD018537) were used in this study.

---

**Table S1.** Species names, accession numbers and collection location are given for samples used to build the phylogeny shown in Figure 1A. Multiple individuals of a species are listed as independent samples with accession numbers and a numerical suffix appended to the species name. Abbreviations: NP (National Park), SP (State Park).

| Sample Name | Accession # | Collection Location |
| --- | --- | --- |
| *Asobara sp. ABZ3773.1* | KR896844.1 | Prince Albert NP, SK, CAN |
| *Asobara sp. ABZ3773.2* | KR898757.1 | Elk Island NP, AB, CAN |
| *Asobara sp. ABZ3773.3* | KR886087.1 | Elk Island NP, AB, CAN |
| *Asobara sp. ABZ3773.4* | KR875914.1 | Prince Albert NP, SK, CAN |
| *Asobara sp. ABZ3773.5* | KR884238.1 | Wellington County, ON, CAN |
| *Asobara sp. ABZ3773.6* | KR888074.1 | Elk Island NP, AB, CAN |
| *Asobara sp. ABZ3773.7* | KR879657.1 | Wellington County, ON, CAN |
| *Asobara sp. ABZ3773.8* | KR784633.1 | Leeds and Grenville, ON, CAN |
| *Asobara sp. ABX5347* | JN293161.1 | Yoho NP, BC, CAN |
| *Asobara sp. ACE4721.1* | JN293665.1 | Glacier NP, BC, CAN |
| *Asobara sp. ACE4721.2* | JN292450.1 | Glacier NP, BC, CAN |
| *Asobara sp. ACR5030* | MF936732.1 | Gulf Islands NP, BC, CAN |
| *Asobara sp. AAE0947* | HQ106668.1 | Restigouche, NB, CAN |
| *Asobara sp. ACF3747.1* | HQ930298.1 | Bigelow, AR, USA |
| *Asobara sp. ACF3747.2* | KR896531.1 | Wellington County, ON, CAN |
| *Asobara sp. ACF3746.01* | HQ929638.1 | Deadhorse Ranch SP, AZ, USA |
| *Asobara sp. ACF3746.02* | KY843128.1 | Islamabad, Pakistan |
| *Asobara sp. ACF3746.03* | KY845150.1 | Islamabad, Pakistan |
| *Asobara sp. ACF3746.04* | KY838943.1 | Islamabad, Pakistan |
| *Asobara sp. ACF3746.05* | KY830675.1 | Islamabad, Pakistan |
| *Asobara sp. ACF3746.06* | KY843285.1 | Islamabad, Pakistan |
| *Asobara sp. ACF3746.07* | KY832326.1 | Islamabad, Pakistan |
| *Asobara sp. ACF3746.08* | KY842379.1 | Islamabad, Pakistan |
| *Asobara sp. ACF3746.09* | KY830249.1 | Islamabad, Pakistan |
| *Asobara sp. ACF3746.10* | KY840132.1 | Islamabad, Pakistan |
| *Asobara sp. AsDen* | MT498809.1 | Denver, CO, USA |

**Table S2.** Species and strain names, accession numbers and collection location are given for samples used to build the phylogeny shown in Figure 1B.

| Species Name | Strain Name | Accession # | Collection Location |
| --- | --- | --- | --- |
| *Asobara sp. ABZ3773* | *ABZ3773* | KR886087.1 | Elk Island NP, AB, CAN |
| *Asobara sp. ABX5347* | *ABX5347* | JN293161.1 | Yoho NP, BC, CAN |
| *Asobara triangulata* | *DSZ062* | KT835413.1 | Yunnan, China |
| *Asobara mesocauda* | *DSZ061* | KT835414.1 | Yunnan, China |
| *Asobara rufescens* | *TK(1)* | AB920758.1 | Tokyo, Japan |
| *Asobara tabida* | *AtFr* | JQ808428.1 | Sospel, France |
| *Asobara citri* | *AcIC* | JQ808423.1 | Lamto, Cote d'Ivoire |
| *Asobara leveri* | *DSZ084* | KT835427.1 | South Korea |
| *Asobara brevicauda* | *DSZ066* | KT835453.1 | South Korea |
| *Asobara persimilis* | *AperAus* | JQ808425.1 | Sydney, Australia |
| *Asobara elongata* | *DSZ048* | KT835452.1 | Yunnan, China |
| *Asobara japonica* | *AjJap* | JQ808424.1 | Tokyo, Japan |
| *Asobara rossica* | *A_rossica* | AB456708.1 | Hokkaido, Japan |
| *Asobara pleuralis* | *Aplndo* | JQ808427.1 | Manado, Indonesia |
| *Asobara unicolorata* | *DSZ055* | KT835410.1 | Yunnan, China |
| *Asobara sp. AsDen* | *AsDen* | MT498809.1 | Denver, CO, USA |

